# Quadruple Parametric Optical Mapping to Measure Cardiac Metabolism-Excitation-Contraction Coupling

**DOI:** 10.1101/2025.03.12.642802

**Authors:** Amber Mills, Sharon Ann George

## Abstract

Cardiac excitation contraction coupling has been studied by recording transmembrane potential (V_m_) and intracellular calcium (Ca^2+^), simultaneously. Recently, imaging metabolic substrates such as NADH in addition to V_m_ and Ca^2+^ has been reported to improve our understanding of cardiac physiology and the interaction between these different aspects of heart function, known as metabolism-excitation-contraction-coupling (MECC). In this study, we developed and validated the quadruple parametric optical mapping technique to simultaneously measure V_m_, Ca^2+^ and two metabolic substrates, NADH and FAD.

Langendorff perfused murine hearts of both sexes were subjected to either pharmacological interventions: 0.1 µM nifedipine or 0.1 µM flecainide, or pathological intervention: ischemia/reperfusion and the four facets of cardiac function were measured. Hearts were stained with RH237 and Rhod2-AM dyes to measure V_m_ and Ca^2+^, respectively. Two LED light sources (520 and 365 nm) were used to excite the dyes and induce NADH/FAD autofluorescence, respectively, and all four optical signals were simultaneously recorded and analyzed to measure12 parameters of cardiac function. Nifedipine significantly altered Ca^2+^ related parameters, including increased Ca^2+^ rise-time and Ca^2+^ decay, and prolonged V_m_-Ca^2+^ delay. Flecainide significantly affected all three aspects of MECC with increased V_m_ rise-time and Ca^2+^ decay, while decreasing conduction velocity (CV) and the redox ratio. Ischemia also altered MECC with increased V_m_ rise-time and NADH intensity, decreased CV and redox ratio.

Using quadruple parametric optical mapping, we demonstrate known cardiac drug response to nifedipine and flecainide and report novel aspects of cardiac function that are modulated by these drugs. These novel drug effects would not have been identified without such a multi-parametric approach to assessing cardiac physiology. Further, we were able to determine chronological changes in MECC during ischemia and reperfusion to better understand the interplay between electrical, mechanical, and metabolic function in the heart.

## Introduction

The origin of cardiac imaging traces back to the early 1900’s when George Mines utilized cinematography to capture the dynamics of a frog heart.^1^ The field of optocardiography, since then, has undergone advancements encompassing 3D printing, panoramic imaging, optogenetics, and multi-parametric optical mapping of the heart. Optocardiography employs light-based methods to study cardiac physiology, offering insights into the dynamics of electrical excitation, that triggers changes in membrane potential, intracellular signaling molecules such as calcium, and metabolic substrates such as NADH and FAD. In the past, transmembrane potential (V_m_) was studied using a glass pipette microelectrode, but this technique has limitations in spatial resolution and requires technical precision.^2,3^ In 1976, Salama and Morad performed the first optical recording of V_m_ in the heart.^4^

Optical mapping began with measurements of V_m_ and over the years, has expanded to include NADH, intracellular ions such as calcium, sodium, potassium, magnesium, and oxidative stress markers.^5–12^ More recently, this technique has been advanced to perform combined measurements of multiple parameters by dual and triple parametric optical mapping.^13–18^ While it is important to study each of these parameters individually in healthy and diseased hearts, complex cardiac dynamics in diseased hearts usually impacts multiple factors of cardiac function and their interactions are missed when only one parameter is measured at a time. For this reason, it has become more popular to simultaneously measure excitation-contraction coupling and more recently, metabolism-excitation-contraction coupling (MECC) in the heart in response to drug treatment and diseases.

Cardiac physiology is remarkably complex, with electrical, mechanical, and metabolic components intricately changing from beat to beat. Electrical activity involves the influx and efflux of ions through ion channels and pumps which result in depolarization and repolarization of the cell membrane. Depolarization of the cell causes the opening of calcium ion channels that trigger a calcium-induced calcium release from the sarcoplasmic reticulum, cytoskeletal binding of these calcium ions and mechanical contraction. These physiological processes produce a high energy demand in the heart that requires a constant supply of ATP.^19^ To properly supply the heart with ATP, mitochondria make up almost half of a cardiomyocyte volume.^20^ If metabolism in these mitochondria is chronically impaired, electrical and contractile functions can be affected and arrhythmias and heart failure can ensue.^21^ Thus, the relationship between metabolism, V_m_, and contraction (MECC) is fundamental to understanding cardiac physiology and is essential to study as a whole.

In this study, we add to the current literature of multiple parameter optical mapping by imaging V_m_, calcium transients, NADH, and flavin adenine dinucleotide (FAD) simultaneously, from the same field of view. Both NADH and FAD are cofactors in cellular respiration and energy metabolism. FAD is a cofactor for metabolic reactions of the flavin protein that utilizes multiple redox states to shuttle electrons.^22^ The resulting electron transfer is important for maintenance of the redox balance and calculating the ratio of FAD to NADH gives the redox ratio of the tissue. All four parameters measured in this technique of quadruple parametric optical mapping are intertwined to affect cardiac function and MECC. When recorded simultaneously the four parameters can give a more complete picture of cardiac physiology, and how they are impacted in states of disease or altered by pharmacology. Here, we report for the first time quadruple parametric optical mapping that uses four cameras to capture NADH, FAD, Ca^2+^, and V_m_ signals at the same time. The addition of FAD allows us to assess the oxidative burden in the heart. Furthermore, FAD is more stable than NADH, providing a consistent recording between animals or during different experimental conditions.^23,24^

## Material & Methods

All procedures were conducted with approval from the University of Pittsburgh’s Institutional Animal Care and Use Committee (IACUC) and in accordance with the NIH Guide for the Care and Use of Laboratory Animals.

### Langendorff Preparation

Male and female mice (∼2 months old) on a C57BL/6 background were used and housed in an AAALAC-certified vivarium facility. Mice were anesthetized, and hearts were quickly excised following cervical dislocation and thoracotomy, and the aorta was cannulated and perfused with a modified Tyrode’s solution (adapted from^11^ (in mM) 130 NaCl, 24 NaHCO_3_, 1.2 NaH_2_PO_4_, 1 MgCl_2_, 5.6 Glucose, 4 KCl, 1.8 CaCl_2_, pH 7.40 and bubbled with carbogen (95% O_2_ and 5% CO_2_) at 37 °C). The heart was allowed to equilibrate in a temperature-controlled (37 °C) bath. Electrocardiogram (ECG) leads were placed in the chamber to record volume conducted pseudo-ECGs, and a pressure transducer was attached to the Langendorf system to measure coronary pressure. The pressure was maintained around 80 mmHg.

A platinum bipolar electrode was placed in the center of the anterior surface of the heart and the cameras were focused onto this surface. Electrical stimuli were applied to determine the threshold of pacing, and once that was determined, hearts were paced at 1.5 times the pacing threshold amplitude and 2 ms stimulus duration during optical recording. The pacing wire was then moved out of the way to allow for the heart to hang unhindered during the perfusion of dyes RH237 (transmembrane potential) and Rhod2-AM (calcium). Dyes were sonicated for 10 minutes in an ultrasonic bath before being injected into the perfusion system. A 5-minute excess dye washout period was introduced after each dye perfusion. To make the dye solutions for injection, a 1 mL syringe was used to mix 30 µL of dye (RH237 at 1.25 mg/ml and Rhod2-AM sat 1 mg/ml) with 970 µL of warmed Tyrode solution (total 1 mL). Then 0.4-0.6 mL was injected into the dye injection port making sure there were no air bubbles in either the port or the syringe. After the dyes were injected, the pacing wire was re-placed, and restitution protocol was applied where the basic cycle length (BCL) of the electrical stimulation was sequentially reduced from 200 ms until loss of 1:1 capture or arrhythmia. The heart was illuminated by two LED excitation light sources at 365 nm and 520 ± 5 nm wavelengths. While the former induces autofluorescence of NADH and FAD in the tissue, the latter excites RH237 and Rhod2-AM dyes. Control optical recordings of NADH, FAD, V_m_, and Ca^2+^ were simultaneously acquired at a 1 kHz sampling rate.

### Pharmacological treatment

Six mice were used (4 males, and 2 females) for this study. In these studies, cardiac motion was arrested by the use of an electromechanical uncoupler, 2,3-butanedionemonoxime (BDM). After initial baseline recordings, hearts were treated with 0.1 μM nifedipine (Cayman, Catalog #11106). After 15-minute delay, optical recordings were acquired as above. Next, nifedipine was washed out by perfusion with control Tyrode’s solution for 10 minutes and restitution protocol was repeated after washout. Finally, the hearts were treated with flecainide (0.1 μM, Millipore Sigma Catalog # F6777). Optical recordings were once again acquired after a 15-minute acclimation period as described above.

### Ischemia – No Flow Ischemia

In a separate set of 5 hearts (2 males, and 3 females), after the initial equilibration period, dye staining/washout, and baseline measurements described above, perfusion to the heart was stopped and the heart was subjected to 5 minutes of no flow ischemia. Optical recordings were obtained at 1-minute intervals during the ischemic period at 150 ms BCL (150 ms BCL is considered normal heart rate for *ex vivo* mouse heart preparation). After the last recording (5 minutes total) perfusion of Tyrode’s solution was reinstated, and optical recordings at 1 min intervals were made during the first 5 minutes of the reperfusion phase of the protocol.

### Data analysis

Optical mapping data of NADH, FAD, Ca^2+^, and V_m_ were analyzed using custom Matlab software, Rhythm 4.0, which will be made available in an open-source format. Rhythm 4.0 is an upgraded version of Rhythm 3.0 used for dual- and triple-parameter optical mapping data analysis.^11^ Using this program, 12 different parameters were measured from the four optical signals. These parameters include, from (V_m_): (1) rise time (V_m_ RT), (2) action potential duration (APD_80_), (3) transverse and (4) longitudinal conduction velocity (CV_T_ and CV_L_), (5) anisotropic ratio (AR); from intracellular calcium: (6) rise time (Ca^2+^ RT), (7) calcium transient duration (CaTD_80_), (8) calcium decay time constant (τ), (9) V_m_-Ca^2+^ delay; from NADH: (10) NADH autofluorescence intensity and from FAD: (11) FAD autofluorescence intensity, and (12) redox ratio.

Upstroke rise time (RT) from V_m_ and Ca^2+^ signals were measured as the time from 20 to 90% of the upstroke of the action potential and calcium transient, respectively. For V_m_, RT is a function of depolarizing currents. For Ca^2+^, RT indicates the time for the influx of calcium into the cell from the extracellular space and release from the sarcoplasmic reticulum. APD_80_ and CaTD_80_ were measured as the time interval between activation time (time of maximum first derivative of the upstroke) and 80% of repolarization and calcium transient decay, respectively. Change in either direction, increase or decrease, in these two parameters can lead to arrhythmia. Conduction velocity (CV) is the speed at which the activation wavefront travels in either the longitudinal (parallel to fiber orientation) or transverse direction (perpendicular to fiber orientation). It is calculated as the ratio of the interpixel distance between two points and the difference in activation time between those two points in the direction of fastest propagation. The ratio of CV_L_ to CV_T_ gives the anisotropic ratio (AR) which is indicative of the ellipticity of the propagating wavefront. As AR increases, the risk of arrhythmia also increases. The Ca^2+^ τ was determined by fitting an exponential to the last 50% (50–100%) of the calcium transient decay phase. V_m_-Ca^2+^ delay was defined as the time interval between the activation time of the V_m_ signal minus the activation time of the Ca^2+^ signal. Increasing the delay between voltage and calcium activation suggests that there is uncoupling between the electrical and mechanical aspects of heart function. NADH intensity was measured as the average of the absolute autofluorescence intensity value in the optical recording. NADH intensity in each heart was normalized to the first measured intensity value in order to avoid interexperimental variability. FAD intensity was measured similarly to NADH except using the FAD camera signal, and the redox ratio (FAD/NADH) was calculated from the non-normalized values of FAD divided by non-normalized values of NADH. The three repolarization parameters, APD_80_, CaTD_80_ and Ca τ, are only reliably measured when motion is arrested in the heart using an electromechanical uncoupler. In the pharmacological treatment studies, BDM was used to arrest motion, but in the ischemia study, BDM was not used and thus these parameters could not be assessed.

### Statistics

All data are reported as mean ± standard deviation (SD) unless otherwise noted. The sample size was 6 hearts for the pharmacological treatment studies and 5 hearts for the ischemia study. An alpha level of 0.05 was used in all tests. Regression analysis was performed on the restitution data using GraphPad Prism Version 10.3.1 software. Nonlinear regression analysis was performed using the least squares regression fitting method. If significance was found in the slope or intercept of the experimental group compared to control or baseline, a one-way ANOVA was used to assess differences in parameters at individual BCLs. Tukey post hoc correction was applied for multiple comparisons.

## Results

### Design and validation of the quadruple parametric optical mapping system

Quadruple-parametric optical mapping system parts were custom-built at the University of Pittsburgh Machine Shop as illustrated in Fig. 1a (labeled schematic) & 1b (CAD model). The tissue bath that housed the mouse heart was 3D printed, while all other parts were machine-built. All metal used was an aluminum tooling plate, which is precision ground on the thickness tolerance, also referred to as Mic 6 plate or jig plate. The paddle to hold the heart and ECG leads in the tissue bath was 3-D printed using Formlabs black resin V5, and the tissue bath was 3-D printed using clear resin V5 both were printed on a Formlabs Form4 printer. All design files for 3D printed hardware, machined metal (in STL format), and data analysis software (Matlab) are available under an open-source license and will be made available. Cameras 1, 2, 3, and 4 were set up to record NADH, FAD, Ca^2+^, and V_m_, respectively. Fig. 1c illustrates the excitation and emission spectra of the dyes and autofluorescent markers used in this study and how they were separated using dichroic mirrors and filters. The measurement of each of the twelve parameters of cardiac physiology measured in this study is illustrated in Figs. 1d and 1e. From V_m_ sensitive dye signals, V_m_ RT, APD_80_, CV_T_, CV_L,_ and AR were calculated. From intracellular calcium indicator signals, Ca^2+^ RT, CaTD_80_, Ca τ, and V_m_-Ca^2+^ delay were measured. As for NADH and FAD intensity, Fig. 1e shows the normalization of each intensity value to the first recording (Control or Baseline at 200 ms BCL) of that individual heart to limit experimental variability.

**Fig. 1:**
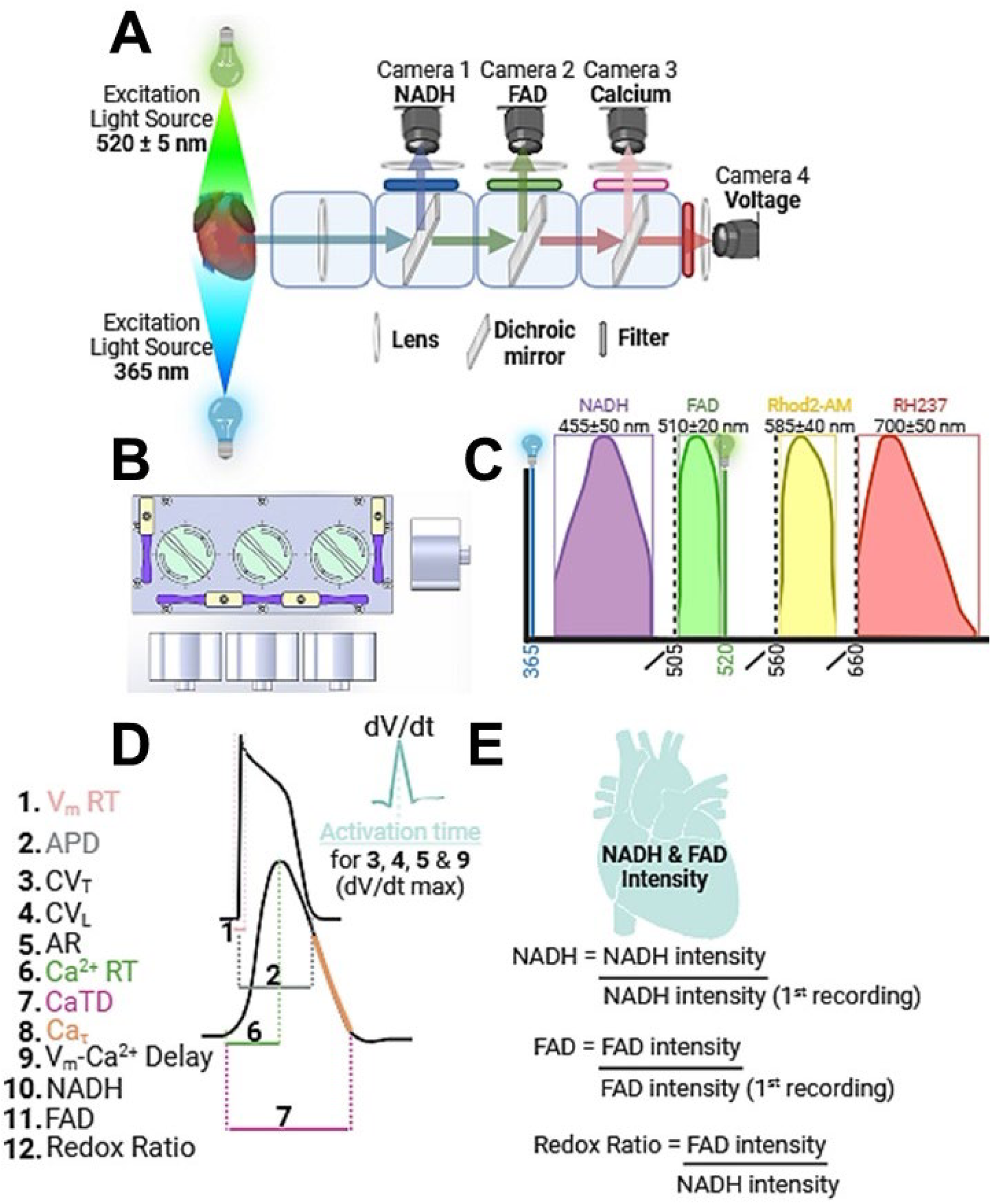
Design and schematic of the quadruple parametric optical mapping system. **A**) Schematic of the system showing a detailed setup for simultaneous optical recordings of NADH, FAD, intracellular Ca^2+^ and V_m_ from the heart. **B**) Computer-aided design (CAD) 2-D illustration of the aluminum machine parts to hold the cameras, lenses, mirrors, and filters. **C**) Spectrum indicating the separation of the four channels to record NADH and FAD autofluorescence and Rhod2-AM (Ca^2+^ indicator dye) and RH 237 (voltage sensitive dye) dyes respectively. The two colored (blue and green) vertical lines represent the 2-LED light source wavelengths, the dotted vertical lines represent the dichroic mirrors used, and the boxes represent the bandpass filters used to ensure 4 unique and separate optical channels. **D**) V_m_, Ca^2+^ activation with color-coordinated explanations of each of the 12 parameters measured during the depolarization and repolarization of the V_m_ signal and calcium release and re-uptake phase from the Ca^2+^ signal. **E**) Normalization equation of NADH and FAD intensity for each heart’s first recording.

As a first step, signals in the four channels were assessed in order to ensure that there was no significant overlap or crossover between the four optical channels. Vials of 1.25 mg/mL RH237 dye, 1 mg/mL Rhod2-AM dye, 500 µg/mL NADH solution, 500 µg/mL FAD solution, and MilliQ water (blank) were placed in the field of view and brought into focus, one at a time. Both the excitation light sources were turned on, and the intensity values within the pixels capturing the vials were recorded. When the NADH solution was imaged, Camera 1 alone recorded a significant increase in intensity compared to Blank (Table 1, p<0.05). Similarly, Camera 2 intensity was increased when FAD solution was imaged, Camera 3 intensity increased when Rhod2-AM was imaged, and Camera 4 intensity increased when RH237 was imaged. Thus, each camera recorded significantly elevated intensities when their respective dye or solution was imaged, and signal overlap was minimal (Table 1, p<0.05).

**Table 1.** Validation of separation of optical channels. Intensity values obtained in the four cameras while imaging of NADH solution, FAD solution, Rhod2-AM (Ca^2+^ indicator dye) or RH237 (voltage sensitive dye) individually during illumination with both excitation light sources are listed below.

| Camera | Reagent Imaged |  |  |  |  |
| --- | --- | --- | --- | --- | --- |
| | Blank | NADH<br>500 $\mu\text{g/mL}$ | FAD<br>500 $\mu\text{g/mL}$ | Rhod2-AM<br>1 $\text{mg/mL}$ | RH237<br>1.25 $\text{mg/mL}$ |
| Cam 1 (NADH) | $194 \pm 402$ | $2527 \pm 421^*$ | $503 \pm 264$ | $732 \pm 149$ | $512 \pm 74.9$ |
| Cam 2 (FAD) | $314 \pm 18.7$ | $413 \pm 112$ | $752 \pm 150^*$ | $229 \pm 93.1$ | $245 \pm 51.2$ |
| Cam 3 (Ca) | $252 \pm 17.9$ | $322 \pm 88.1$ | $406 \pm 159$ | $1241 \pm 289^*$ | $209 \pm 27.1$ |
| Cam 4 ($V_m$) | $205 \pm 3.59$ | $209 \pm 3.87$ | $216 \pm 9.18$ | $230 \pm 23.3$ | $3385 \pm 608^*$ |

### Measuring MECC during pharmacological manipulation with nifedipine and flecainide

Next, we tested the use of the quadruple parametric optical mapping system in determining the modulation of MECC by nifedipine and flecainide. The four optical signals were measured from Langendorff-perfused mouse hearts during control and nifedipine perfusion. Action potentials and calcium transients measured from control and nifedipine conditions are shown in Fig. 2A and representative whole heart maps of V_m_ activation time, Ca^2+^ RT, NADH, and FAD intensities from the same heart before and after nifedipine treatment are shown in Fig. 2B. The modulation of each of the 12 parameters, at varying BCLs, is reported in Fig. 2C. Nifedipine significantly increased Ca^2+^ RT seen in Fig. 2B, and Fig. 2C illustrates that this parameter was elevated at all BCLs with nifedipine compared to control (p=0.0122). Further, V_m_-Ca^2+^ delay was significantly decreased at higher BCLs (100-200 ms) compared to control (p<0.0001). These findings are expected with an L-type calcium channel blocker. Interestingly, nifedipine treatment also increased Ca^2+^ τ, at 150 ms BCL (30.75±4.52 to 37.87±5.89 ms compared to control, p=0.0142). No other measured parameters of MECC were altered by nifedipine treatment at 0.1 μM concentration. Finally, the summary of these results at 150 ms BCL, presented in the 12-parameter panel in Fig. 2D, demonstrates the modulation of only calcium-related parameters with nifedipine, indicating that nifedipine had minimal off-target effects on cardiac function. Ca^2+^ RT and Ca^2+^ τ were increased by 54 and 23% while V_m_-Ca^2+^ delay was decreased by 90% (p<0.0001, p=0.0142, p=0.0124, respectively).

**Fig. 2:**
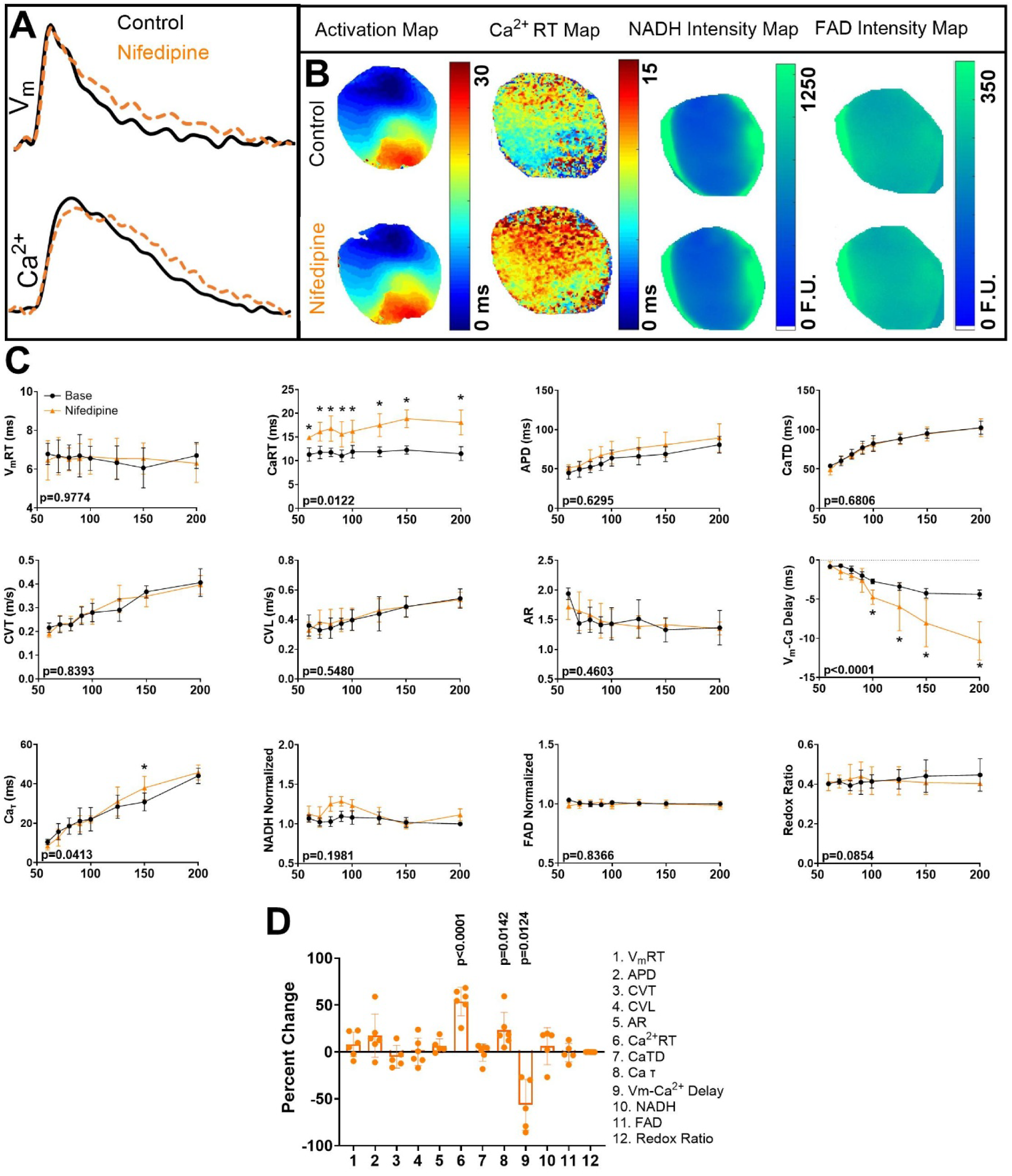
Modulation of cardiac calcium handling by nifedipine. **A**) Representative action potential and calcium transient traces for control (black) and nifedipine-treated (orange) hearts. **B**) Representative whole heart maps showing V_m_ activation, Ca^2+^ RT, and NADH and FAD intensities from control and nifedipine-treated hearts. **C**) Modulation of the twelve parameters of cardiac MECC by nifedipine at varying BCLs. The p-value for slope is listed in the inner corner of each graph determined by linear regression. Only if this was significant (p<0.05) then individual unpaired two tailed t tests was performed. **D**) Summary of the effects of nifedipine on cardiac MECC is illustrated in the twelve-parameter panel indicating the percent change in each parameter versus control at 150 ms BCL. Sample size of 6 hearts was used to generate the data presented in this figure. All significant p-values are reported with Bonferroni correction applied for multiple comparisons.

Next, nifedipine was washed out from these mouse hearts. After confirming the return of all measured parameters to baseline values, these same hearts were exposed to flecainide, a sodium channel blocker. The same 12 parameters of MECC were assessed. Representative action potentials from control and flecainide treatment conditions are illustrated in Fig. 3A while whole heart maps of V_m_ activation time, Ca^2+^ RT, NADH, and FAD intensities are illustrated in Fig. 3B. As expected, flecainide increased V_m_ RT at all BCLs compared to control (p = 0.0074) and decreased both CV_L_ and CV_T_, specifically at the slower pacing rates. APD_80_ was not altered with flecainide treatment. Interestingly, flecainide also modulated Ca^2+^ τ which was increased at 100 and 150 ms BCL compared to the control (Fig. 3C, p<0.05). Lastly, NADH intensity was also significantly increased at the faster pacing rates (60, 70 ms) compared to control (Fig. 3c, p<0.05). No other parameters of cardiac MECC were altered by flecainide treatment. The summary of flecainide-induced modulation of MECC at 150 ms BCL is reported in Fig. 3D. This 12-parameter panel indicated that flecainide modulates all three aspects of cardiac MECC, metabolism, electrical excitation, and calcium handling. Specifically, flecainide increased V_m_ RT and Ca^2+^ τ by 126, and 21%, respectively, while CV_T_ and CV_L_ decreased by 40 and 31% and the redox ratio by 8% (Fig. 3D).

**Fig. 3:**
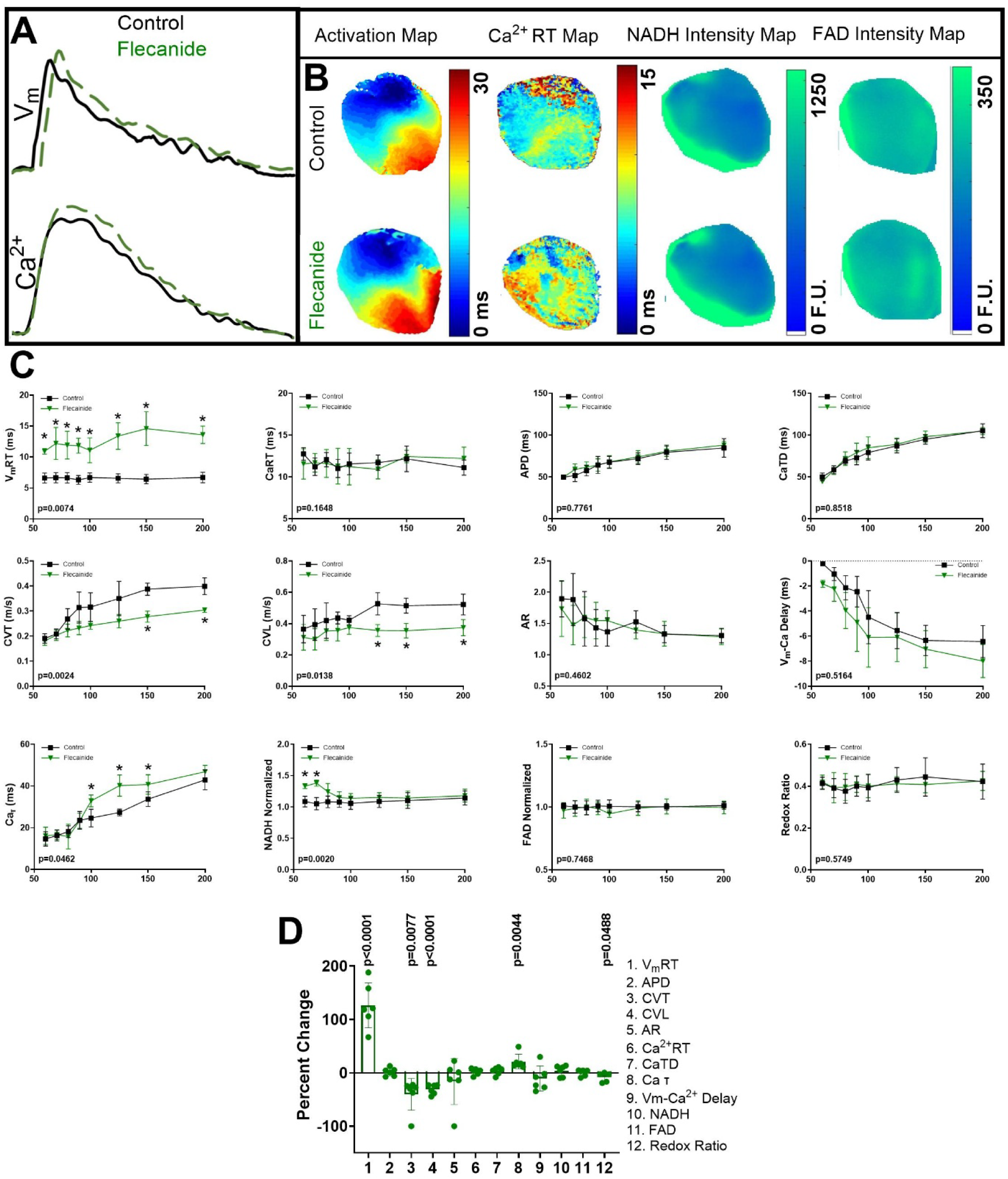
Modulation of cardiac MECC by flecainide. **A**) Representative action potential and calcium transient traces for control (black) and flecainide-treated (green) hearts. **B**) Representative whole heart maps showing V_m_ activation, Ca^2+^ RT, and NADH and FAD intensities from control and flecainide-treated hearts. **C**) Modulation of the twelve parameters of cardiac MECC by flecainide at varying BCLs. The p-value for slope is listed in the inner corner of each graph determined by linear regression. Only if this was significant (p<0.05) then individual unpaired two tailed t tests was performed. **D**) Summary of the effects of flecainide on cardiac MECC is illustrated in the twelve parameter panel indicating the percent change in each parameter versus control at 150 ms BCL. Sample size of 6 hearts were used to generate the data presented in this figure. All significant p-values are reported with Bonferroni correction applied for multiple comparisons.

### Measuring MECC during pathological manipulation with ischemia/reperfusion

Next, a separate set of hearts was subjected to 5 mins of no-flow ischemia followed by 5 mins of reperfusion. During ischemia and reperfusion, optical imaging was performed at 1-minute intervals to determine the sequence of change in MECC parameters. Whole heart maps of V_m_ activation time, Ca^2+^ RT, NADH, and FAD intensities at baseline, ischemia (5 mins), and reperfusion (5 mins) are illustrated in Fig. 4A, and time-dependent modulation of each of the 9 measured parameters are reported in Fig. 4B. First, 5 mins of no-flow ischemia altered both metabolic and electrical excitation parameters in mouse hearts (Fig. 4C). V_m_ RT (4.529 to 7.469 ms) and NADH (1.000 to 1.294) increased within 1 min of ischemia and remained elevated throughout the ischemic period (Fig. 4B). Next, at 3 mins of ischemia, CV_T_ was significantly slower (0.3772 to 0.2162 m/s) and V_m_-Ca^2+^ delay increased (−10.090 to −4.808 ms). These effects also continued until ischemia was reversed. Finally, at 4 mins of ischemia, CV_L_ was also significantly reduced (0.4981 to 0.3092 m/s). When the hearts were reperfused, most of these parameters returned to baseline levels, with the exception of CV_T_ and CV_L_ which increased after 5 mins of reperfusion (16% and 9.2% increase, respectively, Fig. 4C). This overcompensation was also observed with V_m_-Ca^2+^ delay (15% increase, Fig. 4C).

**Fig. 4:**
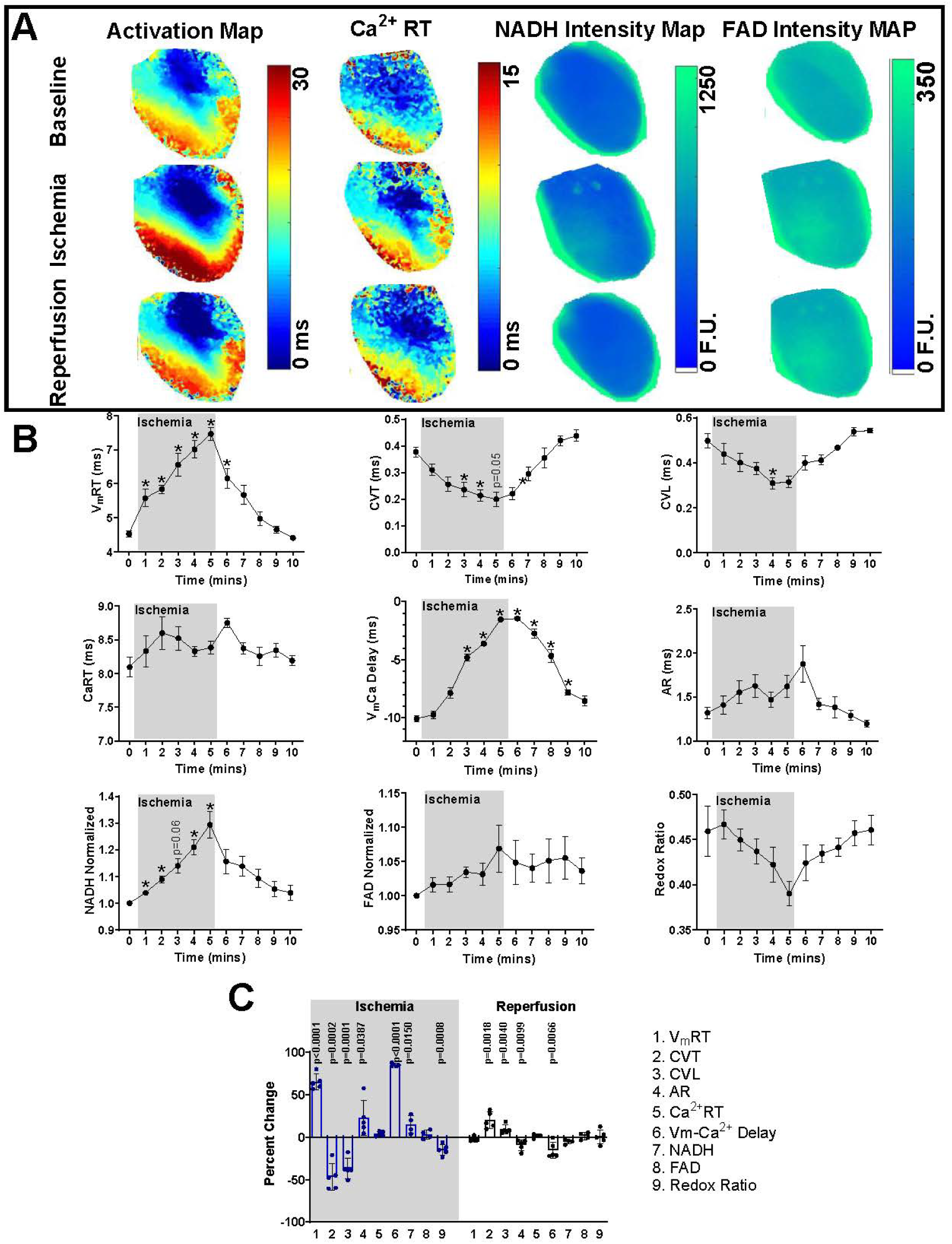
Modulation of cardiac MECC by ischemia and reperfusion. **A**) Representative whole heart maps of V_m_ activation, Ca^2+^ RT, and NADH and FAD intensities during baseline conditions, after 5 mins of no-flow ischemia, and after 5 mins of reperfusion. **B**) Modulation of the nine parameters of cardiac MECC (the three repolarization parameters were not measured due to motion in signals) during 5 mins of ischemia and reperfusion while being paced at 150 ms BCL. The p-value for slope is listed in the inner corner of each graph determined by linear regression. Only if this was significant (p<0.05) then individual unpaired two tailed t tests was performed. **C**) Summary of the effects of ischemia and reperfusion on cardiac MECC is illustrated as the percent change in each measured parameter versus baseline at 150 ms BCL. A sample size of 5 hearts were used to generate the data presented in this figure. All significant p-values are reported with Bonferroni correction applied for multiple comparisons.

## Discussion

We present here the first report of a quadruple parametric optical mapping approach to measure V_m_, intracellular Ca^2+^calcium, NADH and FAD, simultaneously. The open-source system design is provided, and validation of the technique was performed. This approach was then applied to assess MECC with pharmacological and pathological interventions, and 12 parameters of cardiac MECC were measured. We report here the expected as well as novel effects of known compounds nifedipine and flecainide on cardiac MECC and the interaction and sequential modulation of MECC parameters during ischemia and reperfusion.

### Effects of nifedipine on MECC

Nifedipine treatment primarily induced modulation of calcium handling parameters and these included prolongation of Ca^2+^ RT, increased V_m_-Ca^2+^ delay, and Ca^2+^ τ. While the first two effects, increase in Ca^2+^ RT and V_m_-Ca^2+^ delay, are expected with an L-type calcium channel blocker such as nifedipine, its effects on Ca^2+^ τ is a novel finding. Inhibition of the L-type calcium channels slows the influx of extracellular calcium into the cardiomyocyte, which, in turn, results in a longer time for activation of ryanodine receptors and the calcium release from the sarcoplasmic reticulum. Thus, L-type calcium channel block can result in longer time to calcium release after depolarization (increased V_m_-Ca^2+^ delay) and for calcium build up within the cytoplasm (increased Ca^2+^ RT). Increased Ca^2+^ RT can lead to arrhythmias^25^ and if these effects are persistent, cardiac contractions can weaken, and cardiac output can be reduced.^26^ An increase in V_m_-Ca^2+^ delay indicates excitation-contraction uncoupling and disruption of normal cardiac function. Lastly, nifedipine has not been reported to affect calcium reuptake into the sarcoplasmic reticulum or removal from the cytoplasm, however, we report here an increase in Ca^2+^ τ, which indicates that nifedipine treatment increased the time it takes to remove calcium from the cytoplasm at the end of a cardiac cycle. This effect of nifedipine could be due to the modulation of the sarcoplasmic/endoplasmic reticulum Ca^2+^ ATPase 2A (SERCA2A) or sodium calcium exchanger function or it could be due to altered calcium gradients across cellular membranes. These hypotheses were beyond the scope of this technological development and is subject for future investigations. Other than the three calcium-related parameters, nifedipine did not have any off-target effects on cardiac MECC. Nifedipine did not impact voltage parameters, including V_m_ RT, APD, CV_T_, CV_L_ or AR, or metabolic parameters NADH, FAD, and redox ratio. Although it has been previously reported that nifedipine affects APD, this finding is dependent on the concentration used, and the dose of 0.1µm used in this study was not expected to alter APD.^27,28^

### Effects of flecainide on MECC

Contrary to nifedipine, we measured multiple on- and off-target effects of flecainide on cardiac MECC. Firstly, the effects of flecainide, a sodium channel blocker, on V_m_ RT prolongation and CV slowing were expected as previously reported.^29,30^ In addition to these expected on-target effects, we also measured increased Ca^2+^ τ and NADH intensity, particularly at faster pacing rates, and increased redox ratio at 150 ms BCL. While flecainide >5µM concentration has been demonstrated to affect ryanodine receptor function, there have been no reports on the effects of flecainide on calcium reuptake or removal from the cytoplasm.^31^ The mechanism by which flecainide alters calcium decay requires further investigation. Lastly, flecainide also increased NADH levels in the heart and decreased the redox ratio. To the best of our knowledge, this is the first study to report the modulation of cardiac metabolic parameters by flecainide. However, the effects of another sodium channel blocker, lidocaine, and its use in cardioplegic solutions to prevent ischemic injury have been reported.^32,33^ Future investigations into the relationship between sodium channel function and metabolism need to be performed.

### Effects of ischemia and reperfusion on MECC

Ischemia is a complex pathological condition brought on by a lack of oxygenated blood to the heart and is known to affect several MECC parameters.^30^ Even though the effect of ischemia on various MECC parameters is well-established, not all these parameters and their interrelationship are measured simultaneously. Here, we measured V_m_, intracellular Ca^2+^, NADH, and FAD during 5 minutes of no flow ischemia and reperfusion. First, ischemia elevated NADH levels in the heart, within 1 minute of its induction. It has been well documented that ischemia increases NADH because ATP production is reduced.^30,34–37^ Energy is needed for the transfer of NADH and NAD, and with the loss of ATP, NADH levels build.^38^ NADH levels remained elevated and continued to rise throughout the 5 minutes of ischemia with no change in FAD which resulted in a decreasing trend in the redox ratio. Next, ischemia altered multiple parameters related to electrical excitation. Ischemia induced an increase in V_m_ RT, a decrease in V_m_-Ca^2+^ delay, and slowed CV_T_ and CV_L_, as previously reported.^30,39^ Reduced availability of ATP for efficient ion channel function could contribute to these effects. On the other hand, 5 minutes of no flow ischemia and 5 minutes of reperfusion did not affect calcium handling related parameters in this study.

This study is a sequel to our previously published triple parametric optical mapping technique, where we measured electrical excitation, intracellular calcium, and NADH.^30^ In the present study, we additionally recorded changes in FAD levels in the heart in response to pharmacological interventions and ischemia-reperfusion. The simultaneous measurements of both NADH and FAD allowed the calculation of the redox ratio.^30^ While FAD was not significantly altered due to both pharmacological and pathological interventions in this study, the redox ratio was significantly decreased with ischemia (8%) due to NADH accumulation. Previous work testing FAD in an ischemia model found that it was either not altered or slightly decreased.^40,41^ FAD decrease during ischemia is likely due to a lack of oxygen which limits the electron transport chain, preventing the regeneration of oxidized FAD from FADH_2_.^42^ It is for these same reasons that NADH increases with ischemia.^43^ With the loss of oxygen, there is a switch to anaerobic metabolism which produces NADH and mitochondria can no longer oxidize NADH back to NAD.^43,44^ During ischemia, the redox ratio (FAD/NADH) should be reduced, because even if FAD does not change or slightly decreases as we see, NADH is significantly increased.^45^ A decrease in this ratio indicates oxidative stress and is a diagnostic marker to assess the severity of ischemia.^46^ Therefore, being able to measure FAD in addition to NADH is vital to provide a more holistic picture of cardiac function.

Taken together, these data demonstrate the feasibility and necessity of the quadruple parametric optical mapping approach in cardiac pathologies and drug testing studies. Measuring MECC in the heart can improve our understanding of cardiac physiology and thereby improve the development of clinical and pharmacological interventions. Quadruple parametric optical mapping of MECC produced twelve parameters of cardiac function that could be used to study drug toxicity in the heart which is currently limited. Further, this methodology can improve our understanding of cardiac pathological remodeling during an insult such as ischemia and can improve therapeutics.

### Limitations

Due to the nature of the Langendorff system, the heart is removed from all systemic influences such as autonomic and hormonal regulation. There are changes, especially with ischemia and reperfusion, that could impact an *in vivo* heart differently compared to an *ex vivo* preparation. Regardless, this study still demonstrated the capability of the quadruple parametric optical mapping in measuring modulation of cardiac MECC *ex vivo*. Another limitation of the ischemia study in this report is the requirement to not mechanically unload the heart. The use of electromechanical uncouplers such as BDM will reduce the metabolic demand as the heart stops contracting and can provide a less accurate picture of the metabolic parameters measured in this study. As a result of not electromechanically uncoupling the hearts, repolarization and calcium reuptake-related parameters such as APD, CaTD, and Ca^2+^ τ were not measured. To overcome this limitation, we recommend repeating the measurements, first without electromechanical uncoupling and then with it.

## Data availability

All data will be available upon request to the corresponding author.

## Code availability

Rhythm 4.0 will be made available in an open source format upon publication.

## Author contributions

Study conception: A.V.M and S.A.G.; Study design: A.V.M and S.A.G.; Data acquisition, analysis, and interpretation: A.V.M and S.A.G.; Designing and 3D printing hardware: S.A.G.; Preparing analysis software: S.A.G.; Manuscript drafting and figure preparation: A.V.M., and S.A.G.; Revision and final version approval: A.V.M, and S.A.G.

